# Synergistic effects of PARP inhibitors by Schlafen 11 and BRCA2-deficiency through accumulation of single-strand DNA gaps behind a fork

**DOI:** 10.1101/2023.06.28.546820

**Authors:** Hiroshi Onji, Sota Tate, Tomohisa Sakaue, Nobuyuki Onishi, Takashi Matsumoto, Takashi Sugiyama, Shigeki Higashiyama, Junko Murai

## Abstract

Poly (ADP-ribose) polymerase (PARP) inhibitors (PARPis) induce synthetic lethality in breast cancer gene (BRCA)-deficient tumors. Besides the original model proposed by accumulation of double-strand DNA breaks due to the impaired homologous recombination, accumulation of single-strand DNA (ssDNA) gaps due to impaired BRCA-mediated Okazaki fragment processing has emerged as an alternative mechanism of synthetic lethality. Accordingly, PARPis induce ssDNA gaps behind a replication fork in BRCA-deficient cells. Schlafen 11 (SLFN11), a member of the SLFN family, binds replication protein A (RPA)-coated ssDNA gaps and sensitizes cancer cells to DNA-damaging anticancer agents. These facts motivated us to examine the combinational effects of SLFN11 and BRCA-deficiency on PARPis sensitivity. Here, we show that SLFN11 and BRCA2-deficiency synergistically increased sensitivity to PARPis (talazoparib, niraparib, olaparib, and veliparib) at specific concentrations, where SLFN11 alone showed marginal effects. Using chromatin-bound proteins and alkaline BrdU comet assays in human cancer cells, we revealed that ssDNA gaps induced by PARPis were increased by SLFN11 or BRCA2-deficiency and that the combination of the two had the greatest effect. SLFN11 was recruited to and colocalized with chromatin-bound RPA2 under PARPis. SLFN11 recruited around a fork under DNA damage blocked replication, whereas SLFN11 recruited behind a fork under PARPis did not. Chromatin recruitment of SLFN11 and RPA2 was attenuated by the MRE11 inhibitor mirin. Hence, our studies showed that BRCA2-deficiency increased ssDNA gaps behind a fork under PARPis treatment, where SLFN11 bound and further increased the gaps. Our findings provide a mechanistic understanding of favorable responses to PARPis in SLFN11-proficient and BRCA-deficient tumors.

**Significance:** This study reveals how SLFN11 enhances the antitumor effects of PARP inhibitors in BRCA2-deficient cancer cells and highlights the importance of analyzing SLFN11 expression in addition to BRCA analysis in clinical practice.

## Introduction

Poly (ADP-ribose) polymerase (PARP) inhibitors (PARPis) are the first drugs to introduce the concept of synthetic lethality in the clinical setting (1). Initial models suggested that catalytic PARPis prevent the repair of single-stranded DNA (ssDNA) breaks that can turn into double-stranded DNA (dsDNA) breaks when replication forks collide (2,3). Hence PARPis induce synthetic lethality in breast cancer gene (BRCA)-deficient tumors with homologous recombination (HR) defects. Subsequently, clinical PARPis with comparably high catalytic inhibition potency (e.g., olaparib, niraparib, talazoparib, and veliparib) have been developed. One of these inhibitors, olaparib, was first approved by the US Food and Drug Administration in 2014 (4). Based on this initial model, PARPis are selectively toxic to cancer cells harboring *BRCA* mutations or showing HR-deficiency but do not cause toxicity in HR-proficient cells. Although this model has proven applicable when comparing PARPi sensitivity in BRCA-deficient and HR-proficient cells at relatively low concentrations (e.g., 100 nM olaparib) (5,6), PARPis exhibit antitumor effects, even in HR-proficient tumors, in a concentration-dependent manner at relatively high doses (e.g., 1-25 μM olaparib) (4,7,8).

A decade ago, we reported an additional mechanism of action of PARPis, named PARP-trapping (7,8). In the presence of a certain concentration and type of PARPis, PARP1 and PARP2 are trapped at the 5’-DNA ends noncovalently, generating toxic PARP-DNA complexes. Accordingly, PARPis act as “PARP-poisons” because the antitumor effects of PARPis completely disappear in PARP1/2-deficient cells. Importantly, the potency of PARP-trapping is widely different among PARPis, with talazoparib being the strongest, olaparib and niraparib being moderate, and veliparib being the weakest. These differences are reflected in clinical drug dosing; for example, the daily dose of talazoparib is 1 mg, whereas the daily dose of other PARPis is hundreds of mg, indicating that PARP-trapping potency is a limiting factor in deciding the clinical dose (4). Because PARP-trapping is increased by the addition of alkylating agents, e.g., methyl methanesulfonate (MMS) or temozolomide, which generate massive base damage, we proposed that PARP-DNA complexes are generated at the intermediates of base excision repair. Moreover, because olaparib and MMS combination (7) or talazoparib alone (9) block DNA replication, we also proposed that PARP-trapping causes replication blocks and exerts replication stress. Because PARP-DNA complexes lead to the formation of double-strand breaks (DSBs) with bulky PARP proteins upon fork collision, cancer cells need to employ multiple repair factors beyond BRCA to manage PARP-trapping lesions. Recent studies have identified several processing factors for PARP-trapping, such as the metalloprotease SPRTN (10), serine protease FAM111A (11), chromatin remodeling factor ALC1 (12), and p97 ATPase/segregase (13). Thus, PARP-trapping can damage HR-proficient cells but causes more severe damage to HR-deficient cells. Indeed, the clinical indications of PARPis have expanded and are not restricted to cancers with *BRCA* mutations (14). PARP-trapping has been acknowledged as a primary mechanism of action for PARPis (15).

In relation to or regardless of PARP-trapping, recent studies have proposed that the accumulation of ssDNA gaps behind replication forks due to impaired Okazaki fragment processing (OFP) is a primary genotoxic lesion promoting PARPi sensitivity (16–19). However, the mechanisms through which ssDNA gaps are accumulated under PARPis treatment are still controversial, and it is unclear whether PARP-trapping is involved in the impaired OFP. Nevertheless, extensive studies have demonstrated that PARP1 is activated in the process of OFP (16) and that PARP-trapping at unligated Okazaki fragments under PARPis is essential for inducing ssDNA gaps behind a fork (17,19). Furthermore, PARPis have been shown to cause the accumulation of more ssDNA gaps behind a fork in BRCA1- or BRCA2-deficient cells because BRCA1 and BRCA2 are involved in OFP (18). Studies have also shown that the nuclease MRE11 targets stretches of ssDNA gaps behind a fork in BRCA1-deficient cells (17) and that unknown primases (likely PRIMPOL) may fill the ssDNA gaps, which can counteract PARPi sensitivity (20). Notably, the PARP-trapping behind a fork does not greatly affect DNA replication fork rates (19), implying the existence of optimal conditions for PARPis where PARP-trapping causes replication blocks at base damage lesions and/or only causes ssDNA gaps behind a fork.

Schlafen 11 (SLFN11) has become an emerging focus of cancer therapy as it sensitizes cancer cells to a broad range of DNA-damaging anticancer agents causing replication stress (21,22). SLFN11 has an RPA1-binding domain (23) and an ssDNA-binding site (24) and is colocalized with chromatin-bound RPA complexes (23,25). Although the mechanisms of SLFN11-mediated cell killing are not fully understood, one process contributing to cell death is the SLFN11-dependent replication block, which usually occurs within 4 h after drug treatment and continues until cells die (25). SLFN11 is recruited at the replication fork under replication stress because at the stressed fork, the helicase and polymerase are uncoupled, and there is an RPA-coated ssDNA gap in between. SLFN11 binds the gaps on/around forks and somehow blocks replication.

A recent clinical study analyzing the effects of SLFN11 on olaparib sensitivity in patients with ovarian cancer showed that high SLFN11 expression was associated with improved clinical outcomes (26). Intriguingly, subgroup analyses revealed that only patients with *BRCA* mutations benefited from high SLFN11 expression under olaparib treatment. Because SLFN11 functions in the presence of a variety of DNA-damaging agents without regard to BRCA status (9,27), we wondered whether PARPis treatment might affect the underlying mechanisms of SLFN11-dependent toxicity under BRCA-deficient conditions. Moreover, the affinity of SLFN11 to RPA-coated ssDNA and the generation of ssDNA gaps behind a fork by PARPis motivated us to examine the combinational effects of SLFN11 and BRCA-deficiency on PARPis.

In this study, we examined the combinational effects of SLFN11 and BRCA2-deficiency on PARPis sensitivity. Then, we assessed the accumulation of ssDNA gaps under PARPis with SLFN11 and/or BRCA2-deficiency. Our results showed that accumulation was enhanced in the presence of both. Furthermore, SLFN11 colocalized with chromatin-bound RPA2 behind a fork, and this SLFN11 did not block replication but increased ssDNA gaps. Our experiments revealed how SLFN11 enhanced the antitumor effects of PARPis in BRCA2-deficient cancer cells.

## Materials and Methods

### Cell culture and drug treatments

TOV-112D and DAOY cells were purchased from American Type Culture Collection (ATCC; Manassas, VA, USA). Cell line authentication was obtained by short tandem repeat analysis (Promega). Cells were tested negative for mycoplasma contamination. TOV-112D and DAOY cells were grown in Dulbecco’s modified Eagle medium (cat. no. 044-29765; Wako) with 10% fetal bovine serum (cat. no. 536-90165; Hyclone, Cytiva), 20 U/mL penicillin (Meiji-Seika Pharm), and 100 μg/mL streptomycin (Meiji-Seika Pharma) at 37 °C in 5% CO_2_. Camptothecin (CPT) was obtained from Cayman Chemical Company (cat. no. 11694), talazoparib was from MedChemExpress (cat. no. HY-166106), olaparib was from LC Laboratories (cat. no. O-920), and MMS was from Tokyo Chemical Industry (cat. no. M0369). Niraparib (cat. no. S2741), veliparib (cat. no. S1004), and mirin (cat. no. S8096) were obtained from Selleckchem.

### Generation of *SLFN11*-deleted cells

SLFN11-KO cells in DAOY were previously published (28) and SLFN11-KO cells in TOV-112D were generated in another study (Akashi et al., under revision). Disruption of the *SLFN11* gene using the CRISPR/Cas9 method was reported previously (9). Briefly, each guide RNA (5′-gcgttccatggactcaagag-3′ or 5′-gttgagcatcccgtggagat-3′) was inserted into the pX330 plasmid (pX330-SLFN11). Gene-targeting constructs harboring homology arms and a puromycin-resistance cassette were prepared. The targeting construct and pX330-SLFN11 were co-transfected into TOV-112D and DAOY cells by electroporation. After transfection, cells were released into drug-free medium for 48 h followed by puromycin selection until single colonies were formed. Single clones were expanded, and gene-deletion was confirmed by western blotting. This study was approved by Genetic Modification Commission in Ehime University and Keio University and carried out according to the guidelines of the committee.

### Generation of *SLFN11*-overexpressing cells

pPCIP-cSLFN11 was first made by insertion of Xho I-digested 5’- and 3’-terminal inverted repeats cassette sequences of the piggyBac system (29) amplified by PCR using two oligos: 5’– CCGCTCGAGTTAACCCTAGAAAGATAATCATATTGTGACGTACGTTAAAGATAATCATGCGTAAAATTGACGCATGTTCGAAATGCATGG and 5’-CCGCTCGAGTTAACCCTAGAAAGATAGTCTGCGTAAAATTGACGCATGCGAATTCGGTACCATGCATTTCGAACAT GCG into the Sal I-digested pcDNA3 vector (30) to create the pcDNA-IRs. The CAG promoter digested with Mph1103 I and Acc65 I from pCAG-BSD vector (WAKO) cloned into pcDNA-IRs with Mph1103 I and Acc65 I to create pPC. The 3xFL-IRES-PuroR-HSV TK poly (A) signal fragment was PCR amplified from pMX-3xFL-IP (31) using primers 5’-CGGAATTCATGGGCGTTGCCATGCCAGGTGCCGAAGATGATGTGGTGTAACAATTCATGGACTACAAAGACCATG ACGG and 5’-ACATGCATGCGAACAAACGACCCAACACCGTGCGTTTTATTCTGTCTTTTTATTGCCGGTCGACTCAGGCACCGGGCTTGCGGG. The PCR product was cloned into pPC with EcoR I and Pae I to create pPCIP. The full length of SLFN11 cDNA was amplified using primers 5’-AGATCTGGATCCAGCATGGAGGCAAATCAGTGCC and 5’-CACGCGTCTCGAGGCCTAATGGCCACCCCACGGAA, and integrated into the NotI site of the pPCIP vector, creating pPCIP-cSLFN11 vector, using Thermo GeneArt Seamless Cloning and Assembly Enzyme Mix (cat. no. A14606; Thermo Scientific). The final products were validated by sequencing.

The expression vector containing hyperactive PB transposase cDNA under CAG promoter (pCAG2-hyPB) was used before (32). Further plasmid information is available from the Mendeley link (doi: 10.17632/cr443kztj9.1).

An SLFN11 expression vector (pPCIP-cSLFN11) and the expression vector of a hyperactive PB transposase (pCAG2-hyPB) were co-transfected into TOV-112D and DAOY cells by lipofection using Lipofectamine 3000 Reagent (cat. no. L30000-015; Invitrogen, Carlsbad, CA, USA), according to the manufacturer’s instructions. Four days after transfection, cells were incubated in medium containing puromycin (0.2 μg/mL) for another 10 days, and surviving cells were used for the assays.

### Small interfering RNA (siRNA) transfection

Gene-specific siRNAs (mix of 4 sequences) for human BRCA2 (cat. no. L-003462-00-0010) and negative control siRNAs (cat. no. D-001810-10-15) were products of Dharmacon. According to the manufacturer’s instructions, each siRNA (10 nM) was transfected into cells using Lipofectamine RNAimax Reagent (cat. no. 13778-150; Invitrogen). The culture medium was changed 6 h after the transfection, and subsequent experiments were performed 24 h after transfection.

### Antibodies

Antibodies against SLFN11 (cat. no. sc-515071) and CHK1 (cat. no. sc-8408) were obtained from Santa Cruz Biotechnology (Santa Cruz, CA, USA); antibodies against BRCA2 (cat. no. 10741), RPA2 (cat. no. 35869), histone H3 (cat. no. 4499), PAR (cat. no. 83732), PARP1 (cat. no. 9532), phospho-CHK1 (S345) (cat. no. 2348) were from Cell Signaling Technology (Danvers, MA, USA); and antibodies against α-tubulin (cat. no. 017-25031) were from Wako. Human SLFN11 monoclonal antibodies (cat. no. 5-14.12; mAbProtein Co., Ltd., Japan) were used for immunoblotting.

### Immunoblotting and quantification

To prepare whole cell lysates, cells were lysed with Laemmli SDS sample buffer with sonication and boiling. To prepare chromatin-bound subcellular fractions, we followed the protocol for a Subcellular Protein Fractionation Kit for Cultured Cells (cat. no. 78840; Thermo Scientific). Protein concentration in cell lysates were measured using a RC DC Protein Assay Kit (cat. no. 50000119; Bio-Rad Laboratories, Hercules, CA, USA). Immunoblotting was carried out using standard procedures. Secondary antibodies were horseradish peroxidase-conjugated anti-mouse antibodies (cat. no. W402B; Promega, Madison, WI, USA) or rabbit IgG (cat. no. W401B; Promega). Protein signals were visualized using an iBright CL1500 Imaging system (Invitrogen). Quantification of band intensity was performed using ImageJ software.

### Cell viability assay

TOV-112D cells (1.5 × 10^3^) and DAOY cells (1.0 × 10^3^) were seeded into 96-well white plates (cat. no. SPL-30196; SPL Life Sciences) in 100 μL medium per well. Cells were continuously exposed to the indicated drug concentrations for 48 h in triplicate. Cellular viability was determined using an ATPlite 1step Kit (cat. no. 6016731; Perkin Elmer). Briefly, 25 μL ATPlite solution was added to each well of a 96-well plate. After 5 min, luminescence was measured with a FlexStation3 (Molecular Devices) or Varioskan LUX multimode microplate reader (Thermo Scientific). The ATP level in untreated cells was defined as 100%. The viability (%) of treated cells was defined as ATP in treated cells/ATP in untreated cells × 100.

### Apoptosis analysis by flow cytometry

We used an Annexin V-FITC Apoptosis Detection Kit (cat. no. 15342-54, Nacalai Tesque) to detect cell apoptosis, according to the manufacturer’s instructions. Data were collected with a Gallios Flow Cytometer (Beckman Coulter) and analyzed with FlowJo software (Becton Dickinson).

### Immunofluorescence analysis with or without Edu labeling

Cells were seeded on chamber slides (cat. no. 192-004; Watson). If needed, cells were incubated with 10 μM 5-ethynyl-2’-deoxyuridine (EdU) for 1 h before being collected. The cells were pretreated with cold 0.2% TritonX-100/phosphate-buffered saline (PBS) for 1 min on ice. The cells were then fixed with 4% paraformaldehyde in PBS for 15 min and incubated with 3% bovine serum albumin (BSA)/PBS for 10 min. Next, when EdU detection was necessary, cells were incubated with Click-iT Plus reaction cocktail for 30 min (cat. no. C10640; Invitrogen). Cells were incubated overnight with primary antibodies in 3% BSA/PBS at 4 °C. After washing with TBST, the cells were incubated with proper secondary antibodies and Hoechst33342 (cat. no. H3570; Invitrogen) in 3% BSA/TBST for 1 h. Anti-mouse IgG Alexa Fluor 488 (cat. no. A11001; Invitrogen) or anti-rabbit IgG Cyanine3 (cat. no. A10520; Invitrogen) were used as secondary antibodies. After washing with TBST, cells were mounted with ProLong Gold Antifade Mountant (cat. no. P36930; Invitrogen). Images were captured with a Nikon ECLIPSE Ti or ZEISS LSM 900 confocal microscope.

### Data analysis of immunofluorescence microscopy images

The signal intensity in each cell was measured using ImageJ software. A proper-sized circle, slightly larger than the size of a regular cell, was set. The same circle was used to measure the mean intensity of each signal throughout the experiment in isogenic cell lines. The individual signals were plotted. The line plot was obtained using ZEN software. The data were transferred to GraphPad Prism 7 software and illustrated.

### Alkaline BrdU comet assay

We overall followed the previous publication (33). To detect ssDNA gaps, cells were pulse-labeled with 100 µM bromodeoxyuridine (BrdU) (cat. no. 05650-95; Nacalai Tesque) for 1 h, followed by incubation for 90 min as the chase period. We used a Comet Assay Single Cell Gel Electrophoresis Assay kit (cat. no. 4250-050-K; R&D Systems, Bio-Techne) according to the manufacturer’s instructions. Cells (1 × 10^5^/mL) were combined with molten LM Agarose (at 37 °C) at a ratio of 1:10 (v/v) and immediately pipetted onto CometSlides. Plated slides were incubated on a flat surface at 4 °C in the dark for 10 min, immersed in 4 °C lysis solution for 30 min, and then immersed in Alkaline Unwinding Solution for 20 min. Electrophoresis was conducted at 1 V/cm (25 V, 300 mA) for 20 min at 4°C in Alkaline Electrophoresis Solution. Excess electrophoresis solution was drained off, and slides were washed three times for 5 min each by layering with neutralization buffer (0.4 M Tris-HCl, pH 7.4). Slides were then immersed in 70% ethanol for 5 min, dried at 37 °C for 10 min, washed three times for 10 min each in PBS, and blocked with 0.1% Tween20/3% BSA/PBS for 30 min by horizontal layering at room temperature. After blocking, slides were incubated with the mouse monoclonal anti-BrdU antibodies (cat. no. 347580; Beckton Dickinson; 1:250) overnight at 4°C in a humid chamber, then washed three times for 10 min each in TBST and incubated with secondary antibodies (anti-mouse Alexa Fluor 488, cat. no. A11001; Invitrogen; 1:250)/0.1% Tween20/3% BSA/PBS for 1 h at room temperature in the dark. Thereafter, slides were washed three times for 10 min in TBST, covered with coverslips, viewed slides with a Zeiss Axio Observer Z1, and scored with CaspLab software.

### Cell cycle analysis by flow cytometry

Cells were incubated with 10 μM EdU for 1 h using a Click-iT EdU Alexa Fluor 647 Flow Cytometry Assay Kit (cat. no. C10419; Invitrogen) according to the manufacturer’s instructions. Propidium iodide (cat. no. P4170; Sigma-Aldrich, St. Louis, MO, USA) was used to measure DNA content. Data were collected with a Gallios Flow Cytometer (Beckman Coulter) and analyzed with FlowJo software (Becton Dickinson).

### Data availability

The unprocessed and uncompressed imaging data and the proof of cell line authentication are accessible at the following Mendeley link: https://data.mendeley.com/datasets/cr443kztj9/draft?a=235fcf6e-b3be-43d4-ba40-83d475f469ec.

### Statistical analysis

Statistical analyses were carried out using GraphPad Prism 7 software and conducted using one-way analysis of variance with Tukey’s post-hoc multiple comparisons test. Relevant comparisons are shown (NS: not significant, \**P* < 0.05, \*\**P* < 0.01, \*\*\**P* < 0.001, \*\*\*\**P* < 0.0001). Graphs show mean ± standard deviations (SD).

## Results

### SLFN11 enhanced the sensitivity of PARPis in BRCA2-deficient cells

To examine the effects of BRCA2 and SLFN11, we employed the SLFN11-proficient and BRCA2-proficient ovarian endometrioid adenocarcinoma cell line TOV-112D and medulloblastoma cell line DAOY in this study (Figure 1A, S1A). Deleterious mutations in BRCA1 and BRCA2 were not detected in either cell line according to the Cancer Cell Line Encyclopedia database (https://discover.nci.nih.gov/rsconnect/cellminercdb/) (34). The *SLFN11* gene was knocked out using the CRISPR/Cas9 system (Figure 1A, S1A). BRCA2 expression was transcriptionally knocked down using a mixture of siRNAs for *BRCA2* (Figure 1A, S1A). Hereafter, we compare 4 conditions, presented in the following order: SLFN11-knockout (KO)/control siRNA (siCON), SLFN11-KO/BRCA2 siRNA (siBRCA2), parent (SLFN11-proficient)/siCON, and parent/siBRCA2.

**Figure 1.**
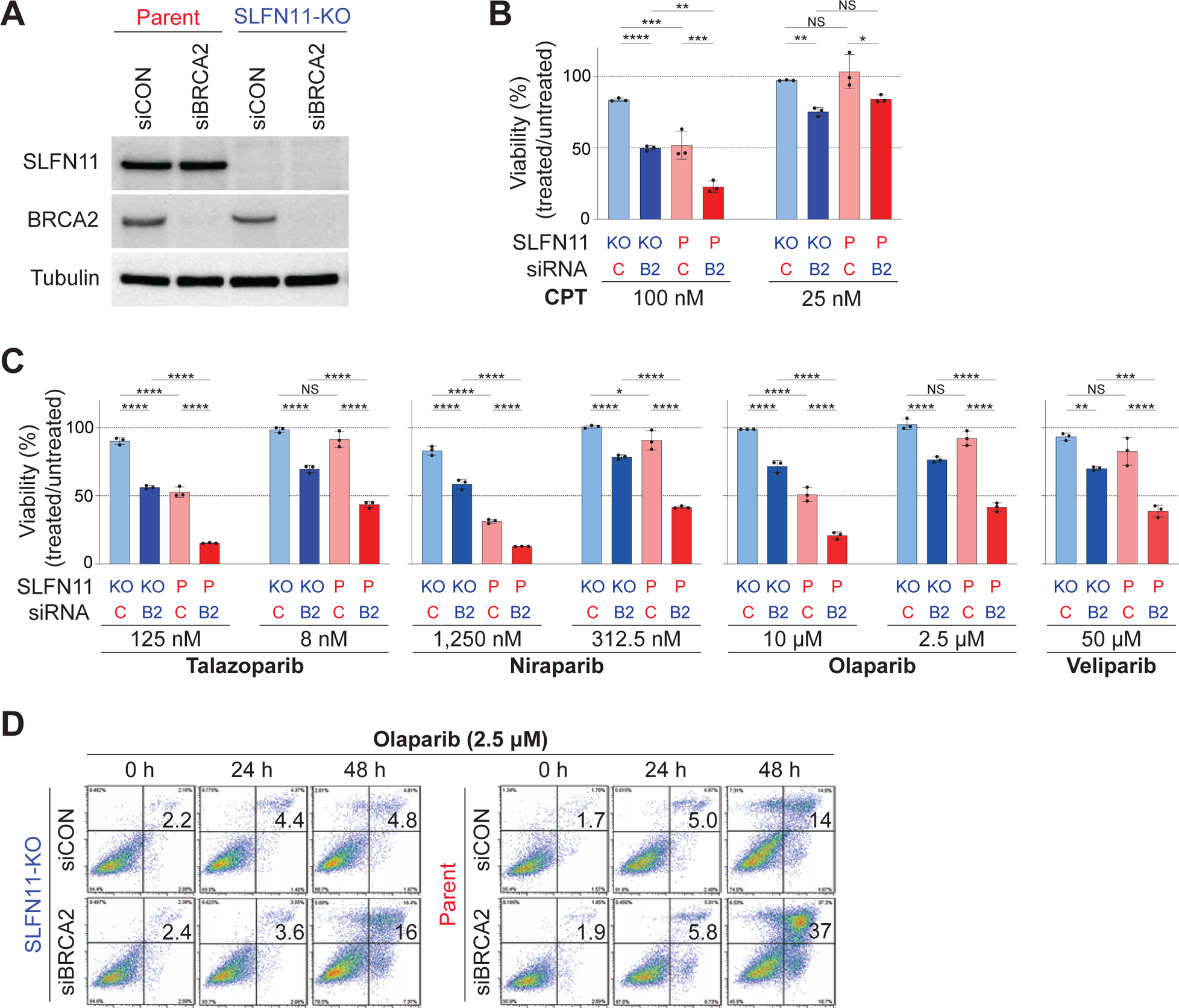
SLFN11 synergistically enhanced sensitivity to PARP inhibitors in BRCA2-deficient cells. **A**, Immunoblot analysis of whole cell lysates prepared from TOV-112D cells: SLFN11-proficient (parent), SLFN11-KO, control siRNA (siCON) or BRCA2 siRNA (siBRCA2). Blots were probed with anti-SLFN11 and anti-BRCA2 antibodies. Tubulin was used as a loading control. **B**, Viability of TOV-112D cells under each condition atier 48 h of continuous CPT treatment. Cellular ATP activity was used to measure cell viability. The viability of untreated cells was set as 100%. Data are means ± standard deviations (n = 3, technical replicates). **C**, Viability of TOV-112D cells under each condition atier 48 h of continuous treatment with PARP inhibitors. Cellular ATP activity was used to measure cell viability. The survival of untreated cells was set as 100%. Data are means ± standard deviations (n = 3, technical replicates). **D**, Apoptosis analysis of TOV-112D cells by flow cytometry treated continuously with 2.5 μM olaparib for 0, 24, or 48 h. NS: not significant, \**P* < 0.05, \*\**P* < 0.01, \*\*\**P* < 0.001, \*\*\*\**P* < 0.0001 (one-way analysis of variance with Tukey’s post-hoc multiple comparisons test).

First, we measured the cellular viability of TOV-112D cells under each condition after 48 h of continuous CPT treatment. Under 100 nM CPT (Figure 1B, left), SLFN11-KO/siCON cells were the most resistant. SLFN11-KO/siBRCA2 cells or parent/siCON cells were more sensitive than SLFN11-KO/siCON cells, and parent/siBRCA2 cells were the most sensitive, suggesting that SLFN11 and BRCA2-knockdown (KD) enhanced the CPT sensitivity additively. Under 25 nM CPT (Figure 1B, right), SLFN11-KO/siBRCA2 cells were more sensitive than SLFN11-KO/siCON cells, but parent/siCON cells were as resistant as SLFN11-KO/siCON cells. Under these conditions, parent/siBRCA2 cells were as sensitive as SLFN11-KO/siBRCA2 cells, suggesting again that SLFN11 and BRCA2-KD enhanced the CPT sensitivity additively. We then examined the effects of BRCA2 and SLFN11 on PARPis (talazoparib, niraparib, olaparib, and veliparib). At relatively high concentrations (125 nM talazoparib, 1,250 nM niraparib, and 10 µM olaparib), we obtained additive effects by BRCA2-KD and SLFN11 in terms of cell viability (Figure 1C). By contrast, at relatively low concentrations (8 nM talazoparib, 312.5 nM niraparib, 2.5 µM olaparib, and 50 µM veliparib), where the effects of SLFN11 were marginal in parent/siCON cells compared with SLFN11-KO/siCON cells, the effects of SLFN11 were significant in parent/siBRCA2 cells compared with SLFN11-KO/siBRCA2 cells. These results implied that SLFN11 could synergistically enhance sensitivity to PARPis under BRCA2-deficient conditions. We obtained comparable results with the DAOY cell set (Figure S1B and C). To further support these findings, we measured the dead cell population under synergistic conditions (2.5 µM olaparib treatment for 48 h) by flow cytometry analysis. Compared with SLFN11-KO/siCON cells, dead cells were increased by 11% and 9% in SLFN11-KO/siBRCA2 and parent/siCON cells, respectively, and by 32% in parent/siBRCA2 cells (Figure 1D). These results demonstrated the synergistic effects of BRCA2-deficiency and SLFN11 on PARPis and suggested the existence of a potential mechanism through which SLFN11 enhanced PARPis sensitivity specifically under BRCA2-deficient conditions.

### SLFN11 and BRCA2-deficiency increased chromatin-bound RPA2 under PARPis

The number of ssDNA gaps is tightly correlated with cell viability under PARPi treatment (18). To examine whether these previous findings were consistent with our results in Figure 1, we measured chromatin-bound RPA2, which reflects the amount of ssDNA in genomic DNA. Immunoblotting demonstrated that chromatin-bound RPA2 tended to increase in TOV-112D SLFN11-KO/siCON cells and increased significantly under the other three conditions in the presence of olaparib (Figure 2A, B). Moreover, chromatin-bound RPA2 was significantly increased in parent/siBRCA2 cells compared with SLFN11-KO/siBRCA2 or parent/siCON cells (Figure 2A, B). Comparable results were obtained with niraparib (Figure S2A). To further support these findings, we measured chromatin-bound RPA2 using immunofluorescence with pre-extraction techniques, which enabled us to specifically detect chromatin-bound proteins. The results matched well with those of immunoblotting (Figure 2C, D). Hence, the amounts of chromatin-bound RPA2 were correlated with cellular sensitivity to PARPis (Figure 1D). By contrast, under CPT treatment, chromatin-bound RPA2 was increased in SLFN11-KO/siCON and SLFN11-KO/siBRCA2 cells at a similar level but was relatively restricted in parent/siCON and parent/siBRCA2 cells (Figure S2B). Hence, the amounts of chromatin-bound RPA2 were not correlated with the viability results (Figure 1B). The results using CPT were consistent with our previous findings demonstrating that SLFN11-proficient cells restrict the extension of uncoupling replication forks that carry ssDNA gaps (25). These results indicated that PARPis were different from CPT in that they increased chromatin-bound RPA2 through the presence of SLFN11 or the absence of BRCA2, with greater effects observed under the combination.

**Figure 2.**
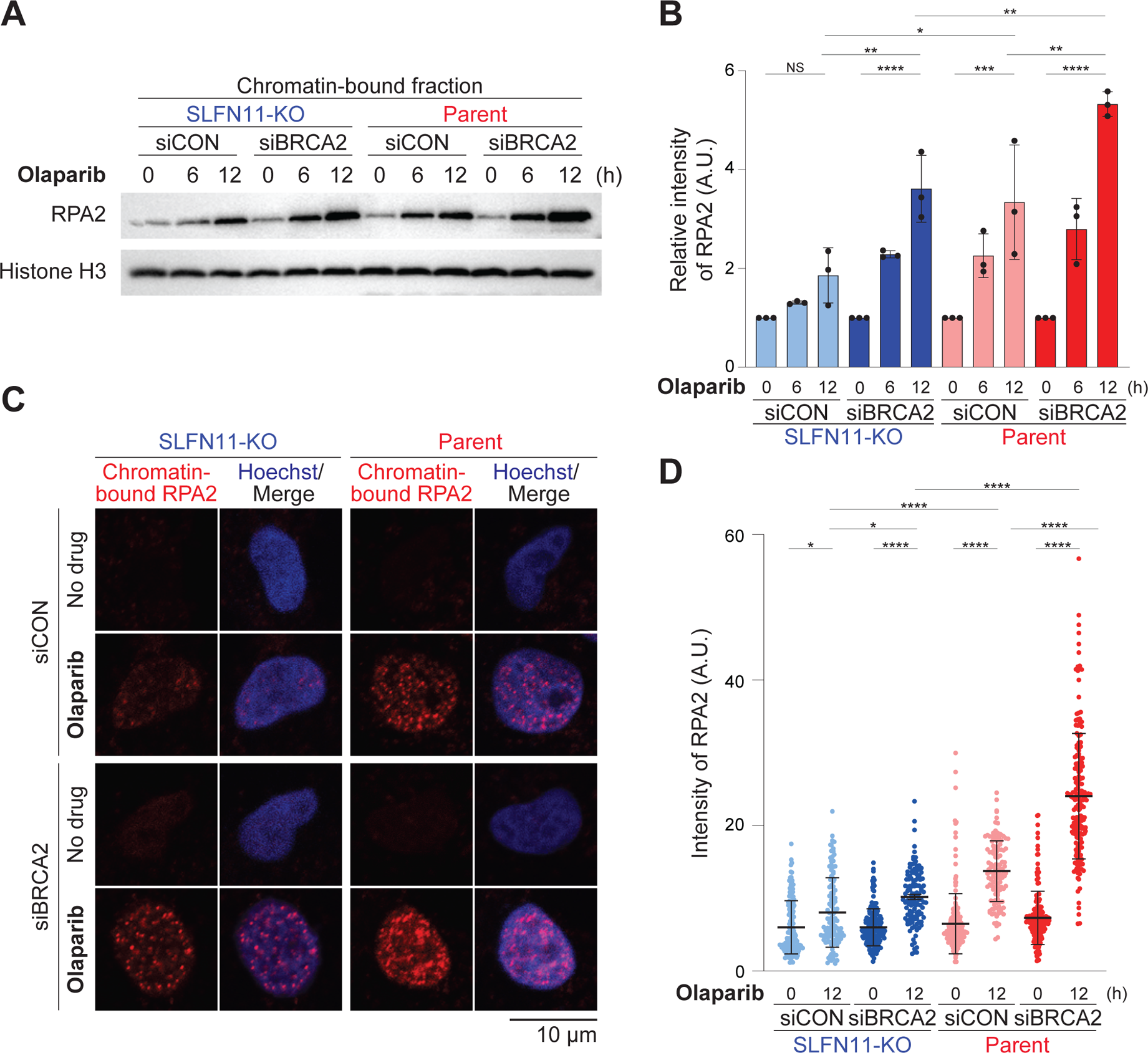
SLFN11 and BRCA2-deficiency increased chromatin-bound RPA2 under PARPis. **A**, Immunoblot analysis of chromatin-bound fractions prepared from TOV-112D cells. Cells were treated with 10 μM olaparib for 0, 6, or 12 h. Blots were probed with anti-RPA2 antibodies. Histone H3 was used as a loading control. **B**, Quantification of data from **A**. Data were normalized to the untreated cells (every 0 h). Data are means ± standard deviations (n = 3, biological replicates). **C**, Representative confocal microscopy images; chromatin-bound RPA2 (red) and Hoechst (blue) in TOV-112D cells. Cells were treated with or without 10 μM olaparib for 12 h. **D**, Quantification of data from **C**. Scatier plots show the mean signal intensity of RPA2. Data are means ± standard deviations (n = 116–199, one-time experiment). NS: not significant, \**P* < 0.05, \*\**P* < 0.01, \*\*\**P* < 0.001, \*\*\*\**P* < 0.0001 (one-way analysis of variance with Tukey’s post-hoc multiple comparisons test).

### SLFN11 and BRCA2-deficiency increased ssDNA gaps in the presence of PARPis

To semiquantify the numbers of ssDNA gaps induced by PARPis, we performed alkaline BrdU comet assays which can be used to analyze the integrity of nascent DNA strands. Under each condition, TOV-112D cells were treated with or without PARPis for 6 h, labeled using BrdU for 30 min during that time, and then incubated in BrdU-free medium for the last 90 min (Figure 3A). After electroporation under alkaline conditions, tail moments of BrdU-labeled cells were measured (Figures 3B, C). Higher tail moment scores can reflect impairment OFP (19). Without drug treatment, tail moments were minimal and were not different among the 4 conditions, suggesting that BRCA2-deficiency or SLFN11 did not impair OFP under normal conditions (Figure 3C). Olaparib (10 µM) treatment significantly increased the tail moment in all conditions but to different degrees (Figure 3B, C). Olaparib potently impaired OFP, even in SLFN11-KO/siCON cells. SLFN11-KO/siBRCA2 cells exhibited significantly higher tail moments than SLFN11-KO/siCON cells, consistent with a previous report showing that BRCA-deficiency impairs OFP (18). Interestingly, parent/siCON cells exhibited significantly higher tail moments than SLFN11-KO/siCON cells, suggesting that SLFN11 also impaired OFP under olaparib treatment. Moreover, parent/siBRCA2 cells exhibited the highest tail moments, possibly reflecting the dual inhibitory effects of SLFN11 and BRCA2-deficiency on OFP. We obtained comparable results with 1.25 µM niraparib treatment (Figure S3).

**Figure 3.**
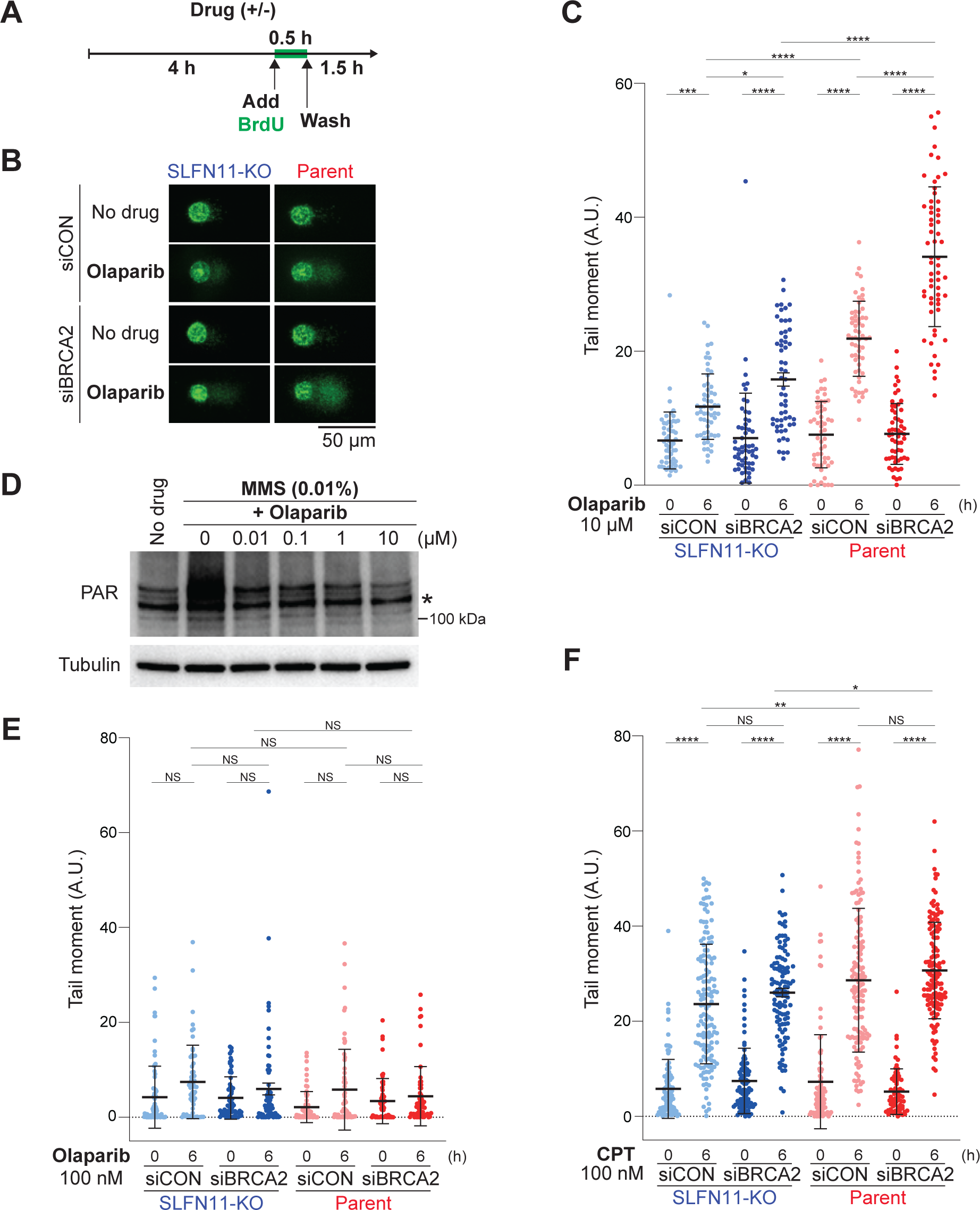
SLFN11 and BRCA2-deficiency increased ssDNA gaps in the presence of PARPis. **A**, Scheme of alkaline BrdU comet assay. TOV-112D cells under each condition were treated with or without drug for 6 h, labeled with BrdU for 30 min during that time, and then incubated in BrdU-free medium for the last 90 min. Atier electroporation under alkaline conditions, tail moments of BrdU-labeled cells were measured. **B**, Representative alkaline BrdU comet assay images in TOV-112D cells treated with or without 10 μM olaparib. **C**, **E**, **F**, scatier plots show BrdU tail moments in TOV-112D cells under each drug treatment (**C**: 10 μM olaparib, **E**: 100 nM olaparib, **F**: 100 nM CPT). Data are means ± standard deviations (**C**: n = 54–59, **E**: n = 54–77, **F**: n = 91–139, one-time experiment). **D**, Immunoblot analysis of PAR levels in whole cell lysates from parent TOV-112D cells treated as indicated for 30 min. Blots were probed with anti-PAR antibodies. The asterisk indicates a nonspecific band. Tubulin was used as a loading control. NS: not significant, \**P* < 0.05, \*\**P* < 0.01, \*\*\**P* < 0.001, \*\*\*\**P* < 0.0001 (one-way analysis of variance with Tukey’s post-hoc multiple comparisons test).

We next examined whether catalytic inhibition or PARP-trapping was more important for OFP impairment. To this end, we first checked the catalytic inhibitory concentration for PARylation by olaparib. Under normal conditions, PARylation was not detectable in TOV-112D cells; however, the alkylating agent MMS increased the PARylation levels in these cells (Figure 3D). The addition of olaparib suppressed PARylation even at 10 nM. Hence, we assumed that 100 nM olaparib would be sufficient to reduce PARylation at a considerably low level. In the presence of 100 nM olaparib, the tail moments were not increased under all conditions (Figure 3E), suggesting that catalytic inhibition of PARP1/2 was not sufficient to impair OFP. By contrast, under 100 nM CPT treatment, the tail moment was increased in all conditions but at similar levels (Figure 3F, 1B). These results confirmed the unique effects of PARPis on impairment of OFP, likely through PARP-trapping. Taken together, these findings demonstrated that olaparib and niraparib impaired OFP and increased ssDNA gaps and that these effects were further enhanced by SLFN11 or BRCA2-deficiency and greatly enhanced by the combination of SLFN11 and BRCA2-deficiency.

### BRCA2-deficiency enhanced the recruitment of SLFN11 on chromatin under PARPis

Given the ability of SLFN11 to bind RPA-coated ssDNA and the finding that olaparib increased the ssDNA gap, we speculated that SLFN11 could be recruited to the ssDNA gap under PARPis. Hence, we collected chromatin-bound fractions from TOV-112D parent/siCON or parent/siBRCA2 cells after olaparib treatment. As expected, olaparib treatment increased chromatin-bound RPA2 in parent/siCON cells and to a greater extent in parent/siBRCA2 cells (Figure 4A, B). Chromatin-bound PARP1 was increased in parent/siBRCA2 cells, indicating that more PARP-trapping lesions were generated under BRCA2-deficient conditions (Figure 4A, B). Chromatin-bound SLFN11 tended to increase in parent/siCON cells and increased significantly in parent/siBRCA2 cells, the kinetics of which were consistent with those of chromatin-bound RPA2 (Figure 4A, B).

**Figure 4.**
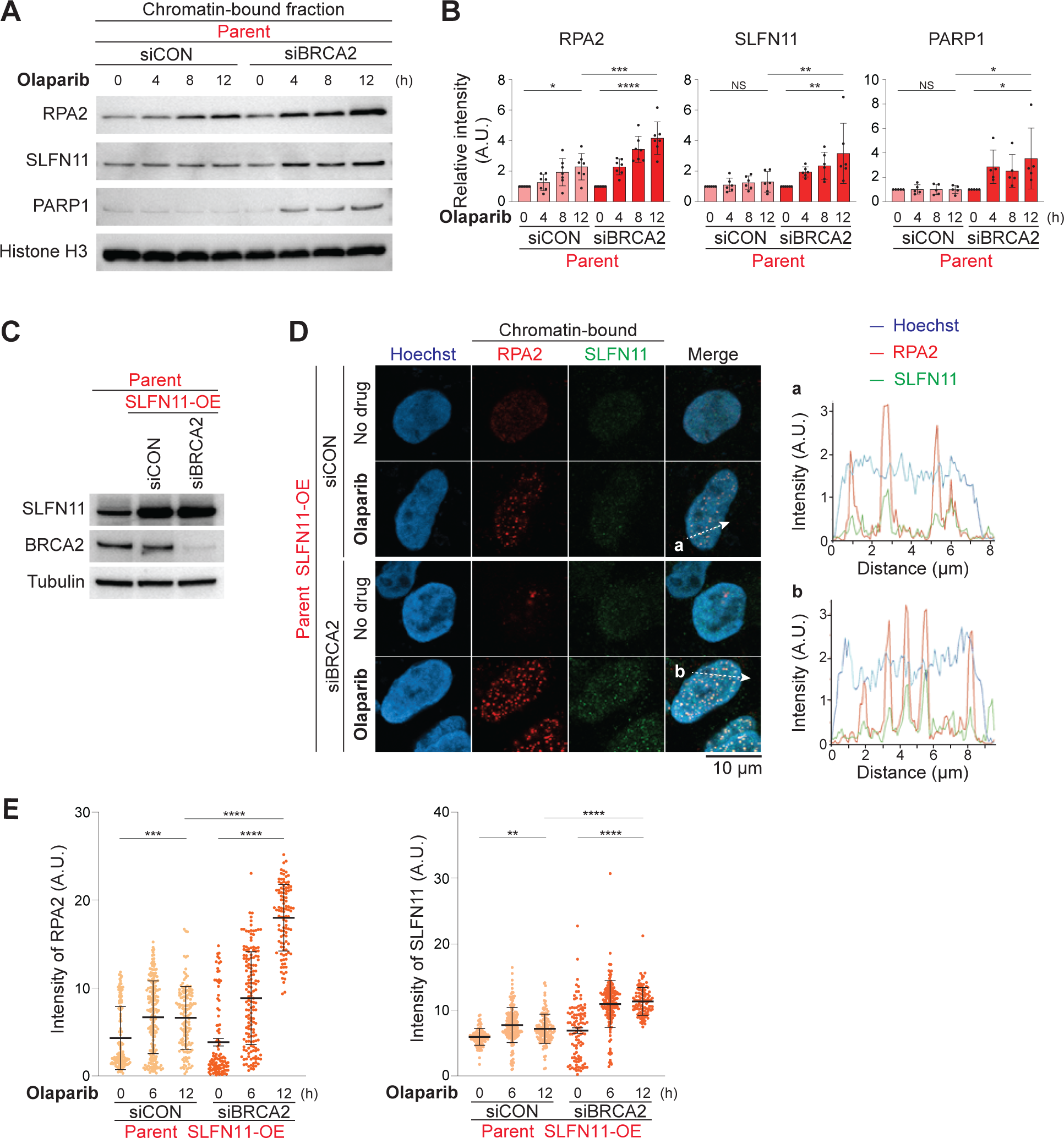
BRCA2-deficiency enhanced the recruitment of SLFN11 on chromatin under PARPis. **A**, Immunoblot analysis of chromatin-bound fractions prepared from TOV-112D cells. Cells were treated with 10 μM olaparib for 0, 4, 8, or 12 h. Blots were probed with anti-RPA2, anti-SLFN11, and anti-PAPR1 antibodies. Histone H3 was used as a loading control. **B**, Quantification of data from **A**. Data were normalized to untreated cells (every 0 h). Data are means ± standard deviations (n = 5–7, biological replicates). **C**, Immunoblot analysis of whole cell lysates prepared from TOV-112D cells. Blots were probed with anti-SLFN11 and anti-BRCA2 antibodies. Tubulin was used as a loading control. **D**, Representative confocal microscopy images; Hoechst (blue), chromatin-bound RPA2 (red), and SLFN11 (green) in TOV-112D cells. Cells were treated with or without 10 μM olaparib for 12 h. Representative tracings of the distribution of signals along the white dashed arrow (**a** and **b**) are shown in the merged panel. **E**, Quantification of data from **D**. Scatier plots show mean signal intensities of RPA2 and SLFN11. Data are means ± standard deviations (n = 104–195, one-time experiment). NS: not significant, \**P* < 0.05, \*\**P* < 0.01, \*\*\**P* < 0.001, \*\*\*\**P* < 0.0001 (one-way analysis of variance with Tukey’s post-hoc multiple comparisons test).

Then, we attempted to detect chromatin-bound SLFN11 using immunofluorescence with pre-extraction. Chromatin-bound SLFN11 was hard to detect because of endogenous SLFN11 expression in TOV-112D parent cells. Therefore, we generated SLFN11-overexpressing (OE) TOV-112D parent cells (Figure 4C). We confirmed that siRNA for BRCA2 in the parent SLFN11-OE cells was as effective as in the original parent cells (Figures 1A and 4C). In parent SLFN11-OE cells, chromatin-bound SLFN11 was increased under olaparib treatment in parent/siCON cells and to a greater extent in parent/siBRCA2 cells (Figure 4D, E). The line plots indicated that chromatin-bound SLFN11 was overall colocalized with chromatin-bound RPA2 (Figure 4D). These results showed that SLFN11 bound to RPA-coated ssDNA gaps under olaparib treatment and that BRCA2-deficiency generated more PARP-trapping and ssDNA gaps behind a fork, where SLFN11 recruitment was enhanced.

### SLFN11 recruited behind a fork did not block replication

We reported that SLFN11 blocked replication on stressed forks under DNA damage. To discriminate the effect of two types of chromatin-bound SLFN11 (around a fork and behind a fork) on replication, we performed cell cycle analyses. In the presence of 100 nM CPT, TOV-112D SLFN11-KO/siCON cells reduced replication at 6 h but resumed replication at 24 h (Figure 5A, top). SLFN11-KO/siBRCA2 cells showed a faster replication state than SLFN11-KO/siCON cells at 6, 12, and 24 h (Figure 5A, second from top). This was likely because the S-phase checkpoint was less activated in SLFN11-KO/siBRCA2 cells than in SLFN11-KO/siCON cells, as supported by the reduced phosphorylation of CHK1 at S345 (Figure S4A) (35). Parent/siCON and parent/siBRCA2 cells reduced replication at 6 h and maintained this replication block at 12 and 24 h (Figure 5A, second from bottom and bottom). The S-phase checkpoint was less activated in parent/siBRCA2 cells than in parent /siCON cells (Figure S4A). However, the effect was not apparent in their replication state, indicating the dominant effects of SLFN11 on the replication state. To examine the chromatin recruitment of SLFN11 under these conditions, we performed immunofluorescence for RPA2 and SLFN11 with pre-extraction and EdU was added 1 h before cell harvesting. Under normal conditions, we did not observe chromatin-bound SLFN11 and RPA2 (Figure 5B, top and second from bottom). After CPT treatment for 12 h, SLFN11 formed foci at the nuclear periphery and the inner nucleus, where RPA2 was colocalized (Figure 5B, second from top and bottom). Cells with SLFN11 foci were mostly negative for EdU signals. At this time point, EdU-positive cells were significantly decreased (Figure 5C, left). These results confirmed our previous findings that SLFN11 around a fork blocks replication.

**Figure 5.**
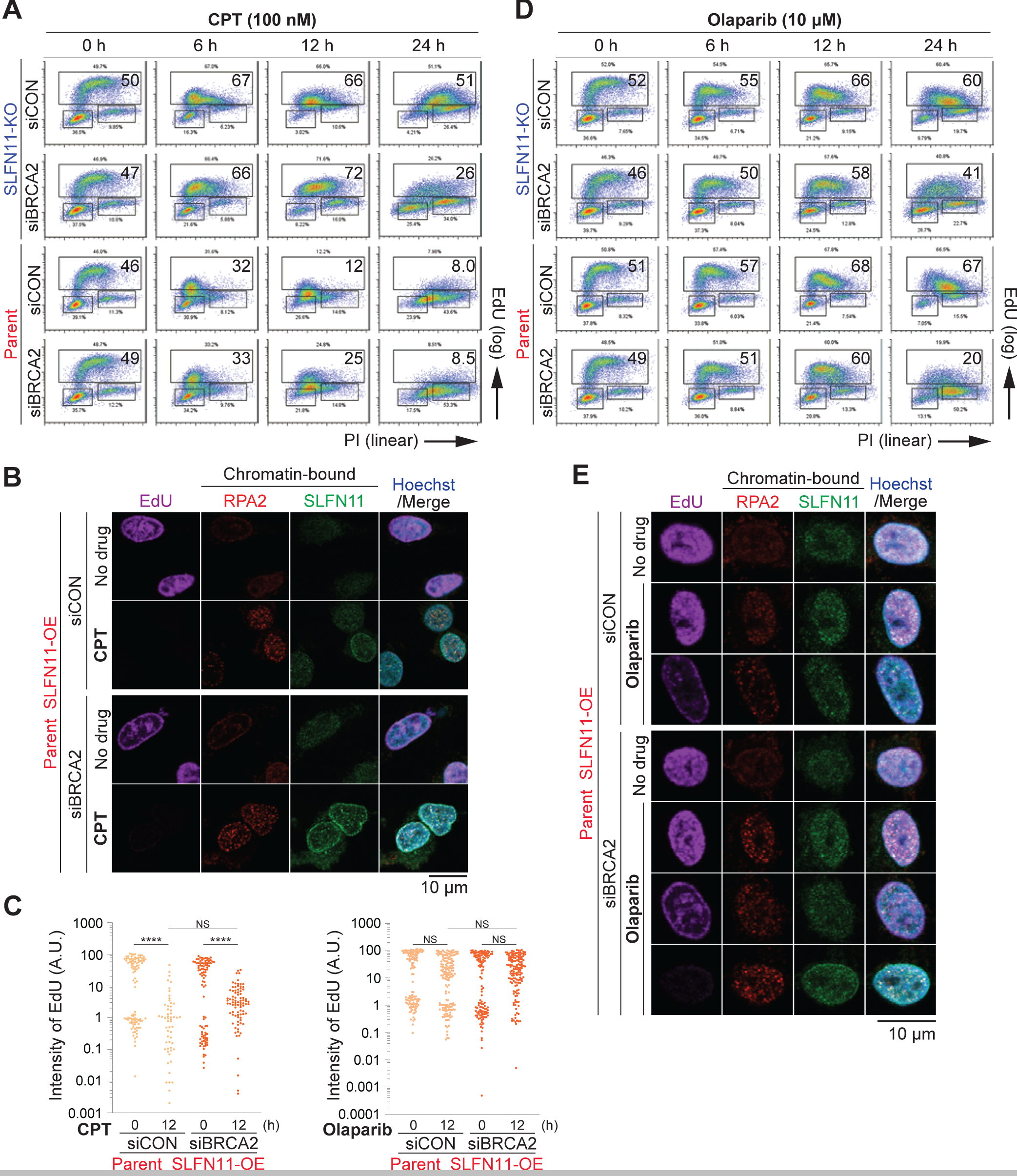
SLFN11 recruited behind a fork did not block replication. **A**, **D**, Representative flow cytometry cell cycle data in response to 100 nM CPT (**A**) and 10 μM olaparib (**D**) atier 0, 6, 12, or 24 h. The percentage of highly replicating cells is annotated. EdU: 5-ethynil-2ʹ-deoxyuridine, PI: propidium iodide. **B**, **E**, Representative confocal microscopy images; EdU (purple), Hoechst (blue), chromatin-bound RPA2 (red), and SLFN11 (green) in TOV-112D cells. Cells were treated with or without 100 nM CPT (**B**) and 10 μM olaparib (**E**) for 12 h. **C**, Quantification of data from **B**, **E**. Scatier plots show the mean signal intensity of EdU. Data are means ± standard deviations (CPT: n = 58–102, olaparib: n = 126–164, one-time experiment). NS: not significant, \*\*\*\**P* < 0.0001 (one-way analysis of variance with Tukey’s post-hoc multiple comparisons test).

We then performed cell cycle analyses in the presence of 10 µM olaparib in TOV-112D cells. SLFN11-KO/siCON cells showed slightly reduced replication at 6 h but continued replicating slowly until 24 h (Figure 5D, top). SLFN11-KO/siBRCA2 cells showed faster replication than SLFN11-KO/siCON cells (Figure 5D, second from top) with less S-phase checkpoint activation (Figure S4B). Parent/siCON cells exhibited cell cycle kinetics similar to those of SLFN11-KO/siCON (Figure 5D, second from bottom and top). Parent/siBRCA2 cells showed faster replication than parent/siCON cells (Figure 5D, bottom) with less S-phase checkpoint activation (Figure S4B). These results indicated that SLFN11 had little impact on the replication state under olaparib treatment. Consistent with these findings, the EdU-positive population was not apparently reduced by olaparib (Figure 5C, right). Chromatin-bound SLFN11 was mostly detected in EdU-positive cells with colocalization with RPA2 under olaparib treatment (Figure 5E). The patterns of SLFN11 foci in the presence of olaparib were clearly different from those in the presence of CPT (Figure 5B and E). Nevertheless, we detected a small number of cells with SLFN11 foci at the nuclear periphery and inner nucleus under olaparib treatment; however, these cells were negatively for EdU overall (Figure 5E, bottom). These results indicated that olaparib had dual effects, namely induction of ssDNA gaps behind a fork and induction of replication stress, both of which recruited SLFN11 to chromatin. However, since cells with SLFN11 foci at a nuclear periphery were a minor population, the former activity seemed to be dominant under olaparib treatment. Overall, comparable results were obtained with the DAOY cell set except that the nuclear periphery pattern of SLFN11 foci was not observed in DAOY parent cells (Figure S4C-F, S5). Taken together, these findings supported that chromatin-bound SLFN11 behind a fork did not block replication.

### Resection by MRE11 was required for SLFN11 to be recruited on chromatin

Given that ssDNA gaps were increased under all conditions, we speculated that the 3’ to 5’ MRE11 exonuclease commonly contributes to the generation of ssDNA gaps under olaparib treatment. To examine the possibility, we treated TOV-112D cells with olaparib and with or without mirin, an MRE11 inhibitor. In SLFN11-KO/siCON and SLFN11-KO/siBRCA2 cells, olaparib treatment increased chromatin-bound RPA2, yet the addition of mirin decreased the RPA2 almost to the control level (Figure 6A, B). In parent/siCON and parent/siBRCA2 cells, increases in chromatin-bound RPA2 and SLFN11 following Olaparib treatment were decreased to the control level by the addition of mirin (Figure 6C, D).

**Figure 6.**
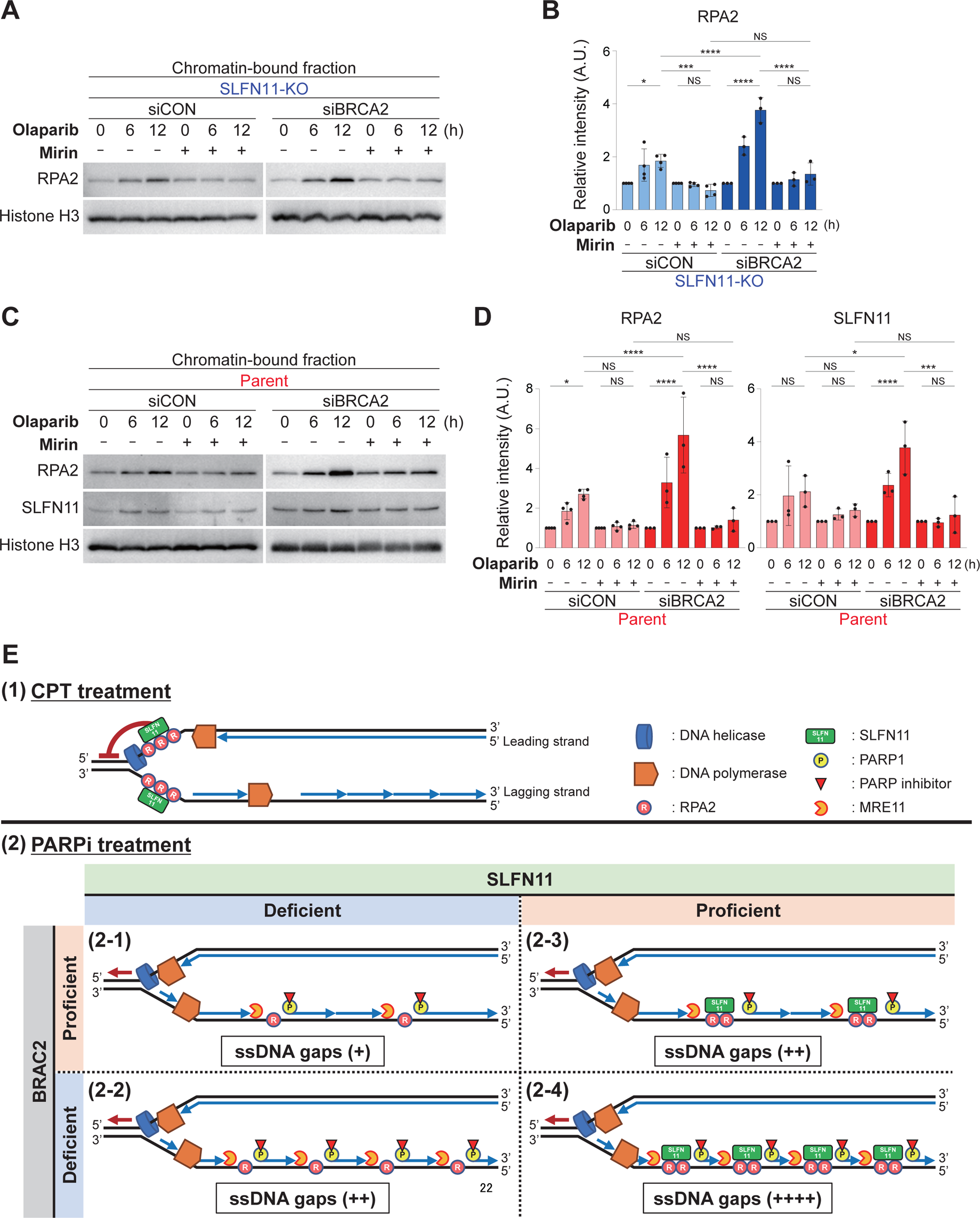
Resection by MRE11 was required for SLFN11 to be recruited on chromatin. **A**, Immunoblot analysis of chromatin-bound fractions prepared from TOV-112D (SLFN11-KO) cells. Cells were treated with 10 μM olaparib and with or without 50 μM mirin for 0, 6, or 12 h. Blots were probed with anti-RPA2 antibodies. Histone H3 was used as a loading control. **B**, Quantification of data from **A**. Data were normalized to untreated cells (every 0 h). Data are means ± standard deviations (n = 3–4, biological replicates). **C**, Immunoblot analysis of chromatin-bound fractions prepared from TOV-112D (parent) cells. Cells were treated with 10 μM olaparib and with or without 50 μM mirin for 0, 6, or 12 h. Blots were probed with anti-RPA2 and anti-SLFN11 antibodies. Histone H3 was used as a loading control. **D**, Quantification of data from **C**. Data were normalized to untreated cells (every 0 h). Data are means ± standard deviations (n = 3–4, biological replicates). **E**, Schematic illustration of the working model. (1) CPT treatment, (2) PARPi treatment. (1) CPT (or a typical DNA damaging agent) exerts replication stress, where the helicase and polymerase become uncoupled, causing the formation of RPA-coated ssDNA gaps in between. SLFN11 binds the gaps on forks and blocks replication. (2-1) In SLFN11-deficient/BRCA2-proficient cells, PARPis generate PARP-trapping, and the DNA strand progressing toward the pre-existing Okazaki fragment is resected by MRE11 due to impaired OFP. The ssDNA gap length or the number of resection sites is limited compared with that under other conditions, and the sensitivity to PARPis is also limited. (2-2) In SLFN11-deficient/BRCA2-deficient cells, ssDNA gaps increase more than in SLFN11-deficient/BRCA2-proficient cells. Moreover, OFP impairment induced by BRCA2-deficiency should contribute to the accumulation of ssDNA gaps. (2-3) In SLFN11-proficient/BRCA2-proficient cells, SLFN11 is recruited to ssDNA gaps atier resection by MRE11 and contributes to the accumulation of ssDNA gaps. (2-4) BRCA2-deficiency causes more ssDNA gaps, as explained in (2-2), where SLFN11 recruitment is enhanced. Consequently, PARPis show the most accumulation of ssDNA gaps in BRCA2-deficient/SLFN11-proficient cells and exhibit the highest anticancer effects. See Discussion for details. NS: not significant, \**P* < 0.05, \*\*\**P* < 0.001, \*\*\*\**P* < 0.0001 (one-way analysis of variance with Tukey’s post-hoc multiple comparisons test).

Hence, resection by MRE11 was likely the first step to generate ssDNA gaps, which occurred regardless of BRCA2 or SLFN11. SLFN11 was recruited to chromatin after the resection by MRE11 and somehow increased the ssDNA gaps.

## Discussion

Accumulation of the ssDNA gaps behind a fork has recently emerged as an alternative mechanism for synthetic lethality in BRCA-deficient cells by PARPis. In this study, we showed that PARPis increased the ssDNA gaps in BRCA2-deficient cells (Figures 2 and 3). Furthermore, we found for the first time that under PARPis treatment, SLFN11 was recruited to RPA-coated ssDNA and increased ssDNA gaps (Figures 2-4). The amount of ssDNA gaps was further enhanced in SLFN11-proficient cells with BRCA-deficiency, which could explain why SLFN11 enhanced the anticancer effects of PARPis in BRCA2-deficient cancer cells (Figure 1). Importantly, 100 nM olaparib was unable to generate the ssDNA gaps (Figure 3), indicating that PARP-trapping was the primary mechanism for inducing ssDNA gaps. Although PARP-trapping, demonstrated by increased PARP1 in the chromatin fraction, was only obvious under BRCA2-deficient conditions (Figure 4A), we assumed that undetectable levels of PARP-trapping should also occur under BRCA2-proficient conditions. Moreover, the increased RPA2 on chromatin was reduced by the addition of mirin under all conditions (Figure 6A-D), indicating that MRE11 functioned to generate ssDNA gaps, regardless of the status of BRCA2 or SLFN11.

Based on these findings, we propose the working model illustrated in Figure 6E. (1) CPT (or a typical DNA damaging agent) exerts replication stress, where the helicase and polymerase become uncoupled, resulting in the formation of RPA-coated ssDNA gaps in between. SLFN11 binds to the gaps on the forks and blocks replication (Figure 5). (2–1) In SLFN11-deficient/BRCA2-proficient cells, PARPis generate PARP-trapping lesions at the 5’-end of Okazaki fragments, and the DNA strand progressing toward the pre-existing Okazaki fragment is resected by MRE11 due to impaired OFP (Figure 6A, B). However, the ssDNA gap length or the number of resection sites is limited compared with those under other conditions (Figures 2 and 3). Hence, PARPi sensitivity is also limited (Figure 1). (2–2) The roles of BRCA2 in the regulation of the S-phase checkpoint have been described only recently (35). Consistent with this, our results showed that BRCA2-deficient cells exhibited faster cell cycle progression than BRCA2-proficient cells likely due to impaired S-phase checkpoint activation (Figure 5B and S4). Hence, the number of Okazaki fragments generated within a certain time duration (e.g., 6 or 12 h) should be higher in BRCA2-deficient cells than in BRCA2-proficient cells. More Okazaki fragments will provide more substrates for PARP-trapping and more resection points for MRE11. Consequently, in SLFN11-deficient/BRCA2-deficient cells, ssDNA gaps increase more than in SLFN11-deficient/BRCA2-proficient cells. Moreover, OFP impairment by BRCA2-deficiency should contribute to the accumulation of ssDNA gaps. (2–3) SLFN11 is recruited to the ssDNA gaps after resection by MRE11 (Figures 6C, D). The mechanisms through which SLFN11 increases ssDNA gaps were not examined in this study. However, chromatin-bound SLFN11 may physically or enzymatically block the function of PRIMPOL, a DNA-directed primase-polymerase that fills the ssDNA gaps (20). Otherwise, SLFN11 on chromatin somehow stabilizes the ssDNA gaps. (2–4) BRCA2-deficiency causes more ssDNA gaps, as explained in (2–2), where SLFN11 recruitment is enhanced. Consequently, PARPis cause the most accumulation of ssDNA gaps in BRCA2-deficient/ SLFN11-proficient cells and exhibit the highest anticancer effect.

The critical point is that SLFN11 around a fork blocks replication, whereas SLFN11 behind a fork does not affect replication but increases ssDNA gaps. The mechanisms of the SLFN11-dependent replication block are only partly understood. However, this study contributes to our understanding of this mechanism by showing that the localization of SLFN11 around a fork should be the key to blocking replication.

### Dual modes of PARPis: PARP-trapping at Okazaki fragments or at intermediates of BER

We have shown that PARP1 and PARP2 are trapped at 5’-ssDNA ends that are the intermediates of base excision repair (BER) arising from endogenous reactive oxidative stress and exogenous base damages (36). Such PARP-trapping lesions are encountered by active replication, resulting in DSBs that cause replication stress. Talazoparib is the strongest PARP-trapping PARP inhibitor and could act in a manner similar to CPT (Figure 6E (1)). Indeed, we have demonstrated the replication block by SLFN11 under talazoparib treatment (9). With talazoparib, PARP-trapping should also occur at Okazaki fragments; however, the replication blocks will suppress the generation of new Okazaki fragments. Thus, the replication block induced by SLFN11 may counteract toxicity from the accumulation of ssDNA. Because a relatively low concentration of talazoparib exhibited synergistic effects in SLFN11-proficient cells with BRCA2-deficiency (Figure 1), the potency or amount of PARP-trapping may determine the mode of action of PARPis. Indeed, the weakest PARP-trapping PARP inhibitor veliparib showed a synergistic effect even at 50 μM (Figure 1). As our findings demonstrated, 10 µM olaparib exhibited dual effects, but the induction of ssDNA gaps behind a fork was dominant (Figure 5). Because the Cmax (maximum concentration in human serum) of olaparib is approximately 20 µM, the dominant mechanism of action for olaparib in the clinical setting will be the generation of ssDNA gaps behind a fork rather than inducing replication block and stress.

These results indicate that PARP-trapping may be more frequent or more easily stabilized at Okazaki fragments rather than BER intermediates (i.e., only strong PARP-trapping inhibitors can generate PARP-trapping at BER intermediates). This can be because the chances of PARP-trapping may be much higher at Okazaki fragments than at BER intermediates. Accordingly, more than 10 million Okazaki fragments must be made and joined in every human cell division (37). By contrast, much fewer BER lesions can be made during replication because an estimated 120,000 base lesions occur in one cell per day (38,39). However, things should be more complex, and we need further studies to clarify these points.

### Implications for the clinical setting

Our findings will provide insights into the mechanisms of favorable responses to olaparib in SLFN11-highly expressing and BRCA-mutant ovarian cancers (26). Hence, our results highlight the importance of analyzing SLFN11 expression in addition to BRCA status in clinical practice. SLFN11 expression level is evaluable with immunohistochemistry in patient tissue samples (40). Importantly, the response of SLFN11 to PARPis is different from that of general DNA-damaging agents (Figure 6E). This notion is important for interpreting clinical data or for when clinical trials are set. Although SLFN11 has other functions, including tRNA-cleavage (41,42), translation impairment (43), chromatin opening (44), degradation of the replication initiation factor CDT1 (45), degradation of reversed replication forks (46), and protection from proteotoxic stress (47), further studies are needed to clarify which function of SLFN11 contributes to the accumulation of ssDNA gaps. Such mechanisms will pave the way for overcoming resistance to PARPis.

### Limitations

In this study, we did not examine the effects of BRCA1. Because Cong et al. reported that ssDNA gaps are increased under BRCA1-deficient conditions (18), we speculate that SLFN11 may also be recruited to ssDNA, and synergize the effects of PARPis in BRCA1-deficient conditions. However, we need to examine whether BRCA1 acts like BRCA2 in terms of regulation of the S-phase checkpoint and whether PARP-trapping and SLFN11 recruitment are increased under BRCA1-deficient conditions.

There is only one clinical dataset analyzing the correlation between SLFN11 expression and response to PARPis with BRCA mutational analyses (26). More clinical evaluations are warranted to further develop our findings for application in the clinical setting.

## Acknowledgments

We thank Mami Chosei and Kenji Kameda from Ehime University and Fumiko Sasaki from Keio University for their great support of our project. This study was supported by the Advanced Research Center (ADRES), Ehime University.

