## Supplemental Figure for "Synergistic effects of PARP inhibitors by Schlafen 11 and BRCA2-deficiency through accumulation of single-strand DNA gaps behind a fork"

### Figure S1

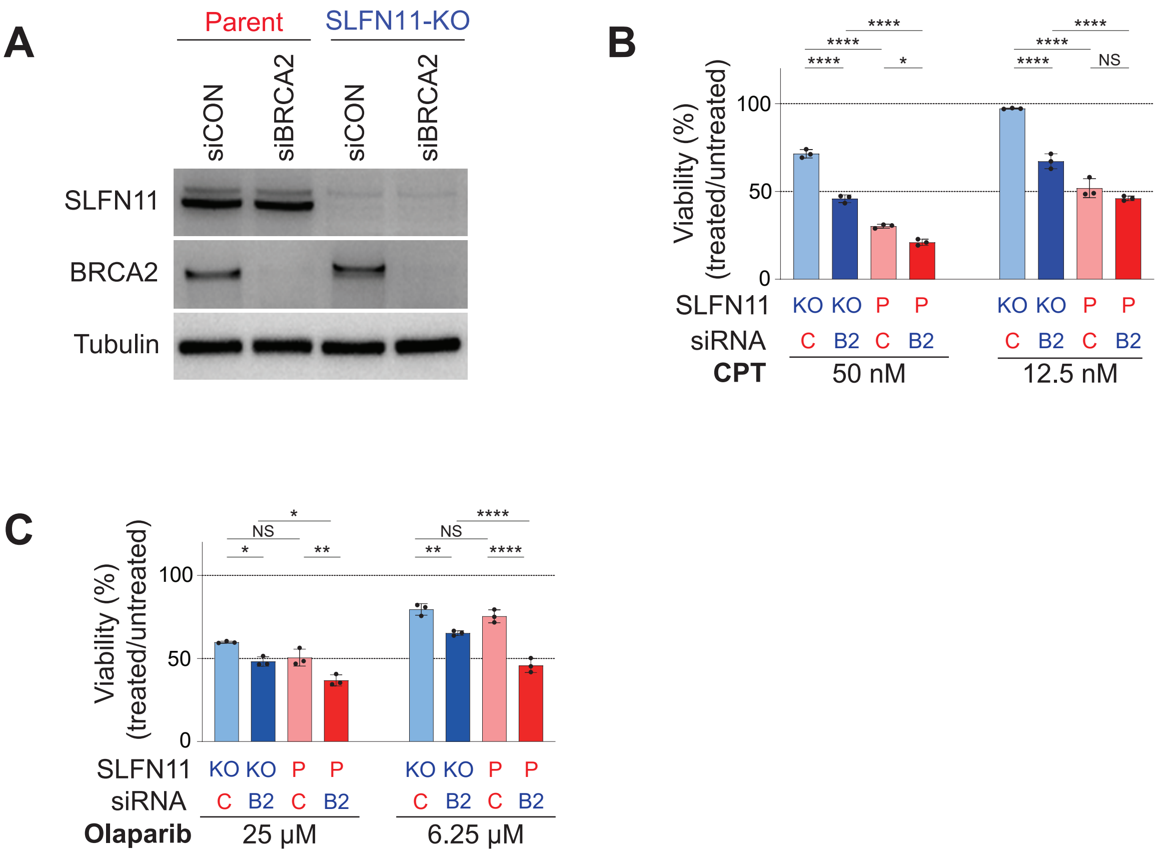


**Figure S1.** SLFN11 synergistically enhanced sensitivity to olaparib in BRCA2-deficient cells. **A**, Immunoblot analysis of whole cell lysates prepared from DAOY cells: SLFN11-proficient (parent), SLFN11-KO, control siRNA (siCON) or BRCA2 siRNA (siBRCA2). Blots were probed with anti-SLFN11 and anti-BRCA2 antibodies. Tubulin was used as a loading control. **B**, **C**, Viability of DAOY cells under each condition after 48 h of continuous treatment with CPT and olaparib. Cellular ATP activity was used to measure cell viability. The survival of untreated cells was set as 100%. Data are means ± standard deviations (n = 3, biological replicates). NS: not significant, *$P < 0.05$, **$P < 0.01$, ****$P < 0.0001$ (one-way analysis of variance with Tukey's post-hoc multiple comparisons test).

Figure S2

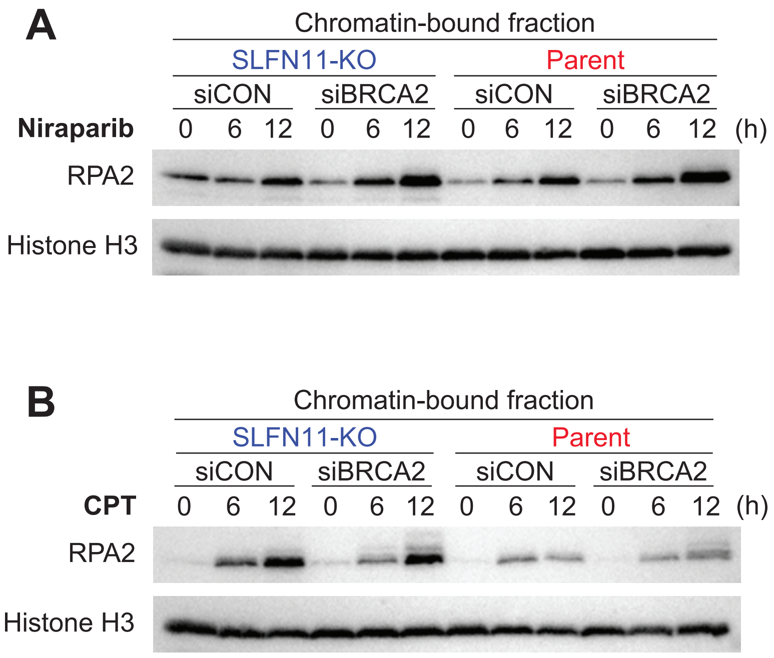


**Figure S2.** SLFN11 and BRCA2-deficiency increased chromatin-bound RPA2 under PARPis. **A**, **B**, Immunoblot analysis of chromatin-bound fractions prepared from TOV-112D cells. Cells were treated with 1.25 μM niraparib and 100 nM CPT for 0, 6, or 12 h. Blots were probed with anti-RPA2 antibodies. Histone H3 was used as a loading control.

### Figure S3

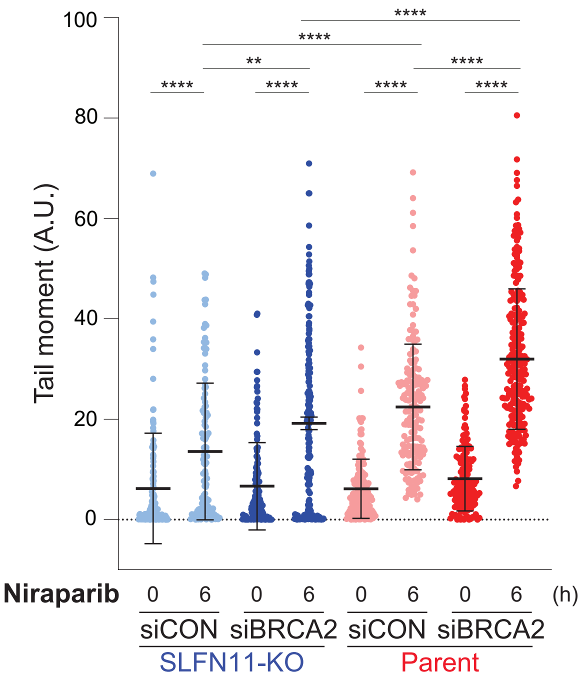


**Figure S3.** SLFN11 and BRCA2-deficiency increased ssDNA gaps in the presence of niraparib. Scatter plots show BrdU tail moments in TOV-112D cells under niraparib 1.25 μM treatment. Data are means ± standard deviations (n = 119–241, one-time experiment). ***P* < 0.01, *****P* < 0.0001 (one-way analysis of variance with Tukey's post-hoc multiple comparisons test).

Figure S4

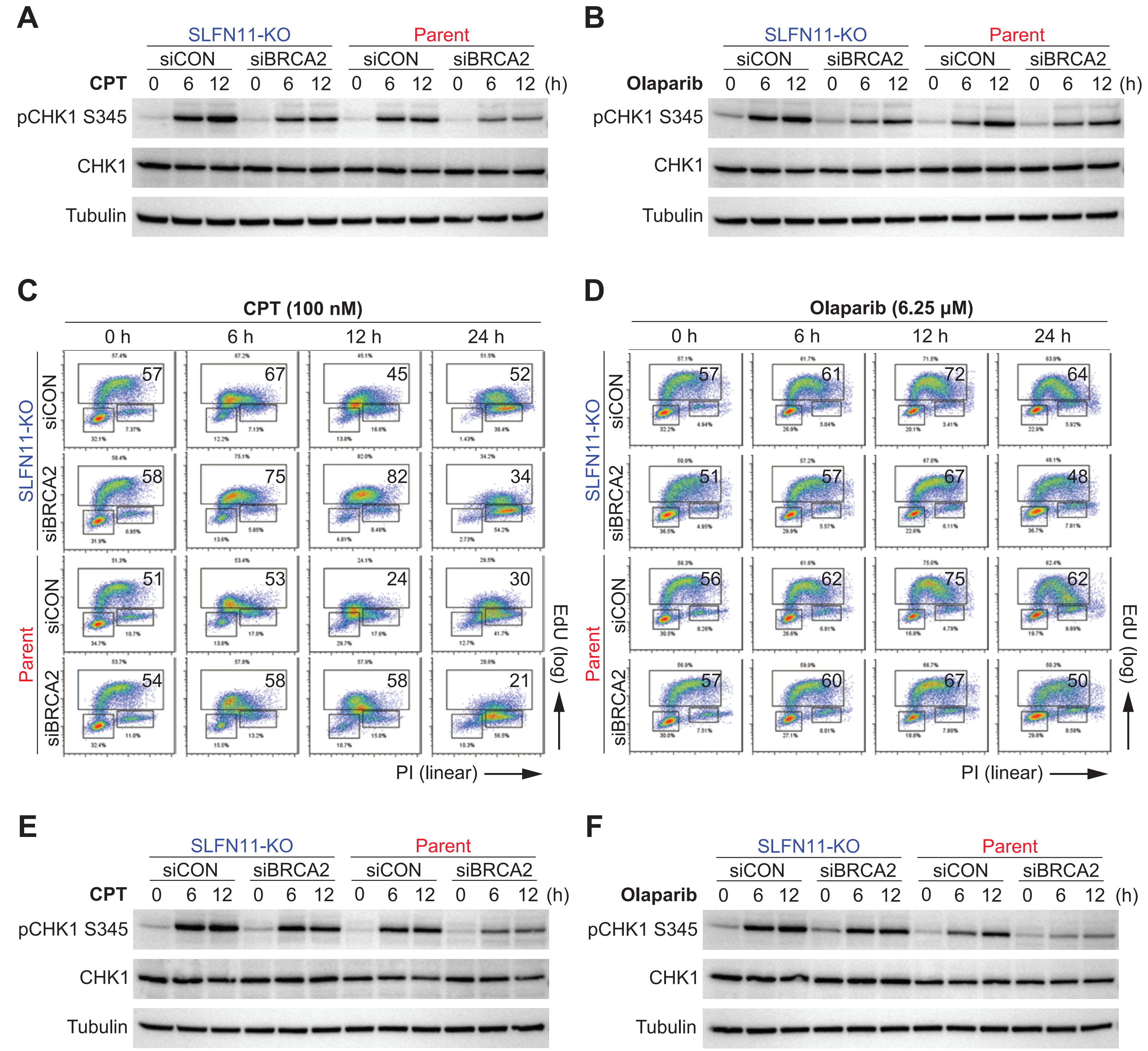


**Figure S4.** The S-phase checkpoint showed less activation in siBRCA2 cells than in siCON cells, which was supported by reduced phosphorylation of CHK1 at S345. SLFN11 recruited behind a fork did not block replication. **A**, **B**, Immunoblot analysis in whole cell lysates from TOV-112D cells treated with 100 nM CPT (**A**) and 10 μM olaparib (**B**). Blots were probed with anti-phospho-CHK1 (S345) and anti-CHK1 antibodies. Tubulin was used as a loading control. **C**, **D**, Representative flow cytometry cell cycle data in response to 100 nM CPT (**C**) and 6.25 μM olaparib (**D**) after 0, 6, 12, or 24 h. The percentage of highly replicating cells is annotated. **E**, **F**, Immunoblot analysis in whole cell lysates from DAOY cells treated with 100 nM CPT (**E**) and 6.25 μM olaparib (**F**). Blots were probed with anti-phospho-CHK1 (S345) and anti-CHK1 antibodies. Tubulin was used as a loading control.

Figure S5

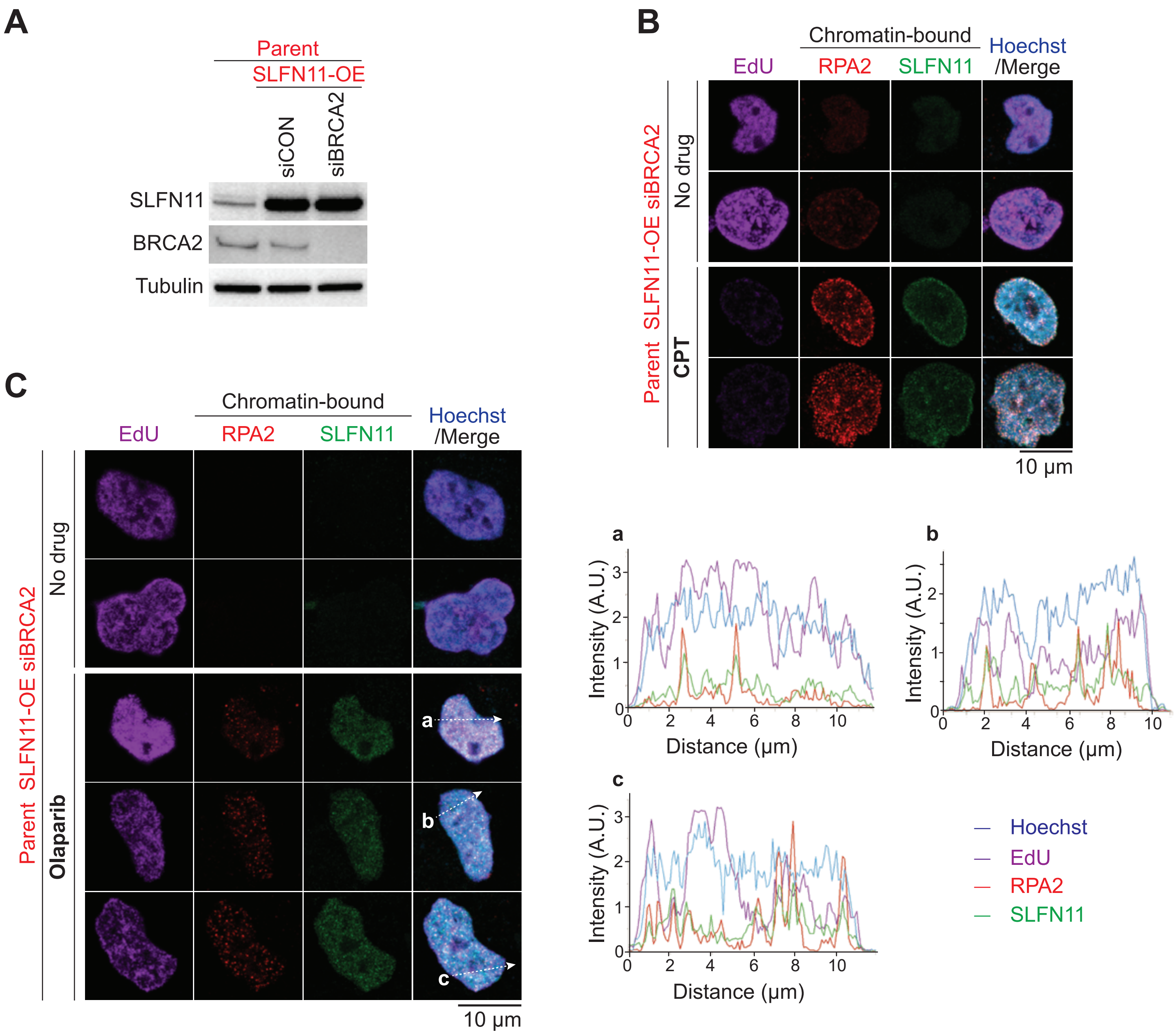


**Figure S5.** SLFN11 recruited behind a fork does not block replication. **A**, Immunoblot analysis of whole cell lysates prepared from DAOY cells. Blots were probed with anti-SLFN11 and anti-BRCA2 antibodies. Tubulin was used as a loading control. **B**, **C**, Representative confocal microscopy images; EdU (purple), Hoechst (blue), chromatin-bound RPA2 (red), and SLFN11 (green) in DAOY cells. Cells were treated with or without 100 nM CPT for 6 h (**B**) and 10 µM olaparib for 12 h (**C**). Representative tracings of the distribution of signals along the white dashed arrow (**a**, **b**, and **c**) are shown in the merged panel.
